# Intronic miR-6741-3p is involved in oral squamous cell carcinoma pathogenesis by targeting the oncogene *SRSF3*

**DOI:** 10.1101/2023.12.20.572638

**Authors:** Dhanashree Anil More, Nivedita Singh, Radha Mishra, Harsha Pulakkat Muralidharan, Kodaganur Srinivas Gopinath, Champaka Gopal, Arun Kumar

**Author notes:** Corresponding author: (AK).

## Abstract

Epigenetic silencing through methylation is one of the major mechanisms for downregulation of tumor suppressor miRNAs in various malignancies. The aim of this study was to identify novel tumor suppressor miRNAs which are silenced by DNA hypermethylation and investigate the role of at least one of these in oral squamous cell carcinoma (OSCC) pathogenesis.

We treated cells from an OSCC cell line SCC131 with 5-Azacytidine, a DNA methyltransferase inhibitor, to reactivate tumor suppressor miRNA genes silenced/downregulated due to DNA methylation. At 5-day post-treatment, total RNA was isolated from the 5-Azacytidine and vehicle control-treated cells. The expression of 2,459 mature miRNAs was analysed between 5-Azacytidine and control-treated OSCC cells by the microRNA microarray analysis.

Of the 50 miRNAs which were found to be upregulated following 5-Azacytidine treatment, we decided to work with miR-6741-3p in details for further analysis, as it showed a mean fold expression of >4.0. The results of qRT-PCR, Western blotting, and dual-luciferase reporter assay indicated that miR-6741-3p directly targets the oncogene *SRSF3* at the translational level only. The tumor-suppressive role of miR-6741-3p was established by various *in vitro* and *in vivo* assays.

Our results revealed that miR-6741-3p plays a tumor-suppressive role in OSCC pathogenesis, in part, by directly regulating *SRSF3*. Based on our observations, we propose that miR-6741-3p may serve as a potential biological target in tumor diagnostics, prognostic evaluation, and treatment of OSCC and perhaps other malignancies.

## Introduction

Oral squamous cell carcinoma (OSCC), arising from surface epithelium, accounts for more than 90% of all oral malignancies [1]. Tumors arising in the oral and oropharyngeal mucosa, including those of the tongue and lips collectively represent the 16^th^ most common cancer worldwide with an annual incidence of 377,713 new cases and 177,757 deaths [2,3]. In India, oral cancer is the most common cancer in males and the fourth most common cancer in females with an ASR (age-standardized rate) of 14.8 and 4.6 per 100,000 males and females respectively [2,3]. Despite recent advances in cancer diagnosis and treatment, including targeted therapy against EGFR using the monoclonal antibody Cetuximab, the 5-year survival rate for oral cancer has remained less than 50% over the last 50 years [4,5]. The current scenario, therefore, requires the identification of new therapeutic targets and molecular markers to aid in better prognosis and treatment.

In recent years, the critical role of microRNAs (miRNAs/miRs) in cancer progression has been widely acknowledged. MicroRNAs belong to a class of endogenous small non-coding RNA molecules of 20-25 nucleotides length, found mostly in eukaryotes, and regulate gene expression by inducing degradation or inhibiting translation of target mRNAs by binding to either the 3’UTRs (untranslated regions), the 5’UTRs or the coding sequences (CDSs) of their target mRNAs [6]. Apart from their specific functions in biological processes such as cell proliferation, differentiation, apoptosis, development, etc., miRNAs are dysregulated in a wide variety of diseases such as immune disorders, Alzheimer’s disease, cardiovascular diseases, rheumatoid arthritis, cancer, etc. [7]. Many functional studies and clinical analysis have linked miRNA dysregulation as a causal factor for cancer progression [8]. Numerous studies have shown that microRNA-based therapeutics hold promising potential for cancer management and hence there is a growing need to further explore their roles in various cancers with the aim of developing miRNA therapeutics [7,8].

MicroRNAs can function as oncogenes or tumor suppressor genes and drive the process of carcinogenesis. The involvement of several tumor suppressor miRNAs like miR-15a, miR-16a, let-7 family, miR-143, miR-145, and oncogenic miRNAs like miR-155, miR-17 cluster and miR-21 in several malignancies including OSCC has been established [8,9]. Aberrant expression of tumor suppressor and oncogenic miRNAs drives the progression from oral premalignant lesions to cancer and also correlates with and could explain the pathogenesis, metastasis, and chemoresistance of OSCC [10,11]. Of the various mechanisms reported to be involved in miRNA deregulation such as loss or mutation of miRNA-encoding genes, defective biogenesis pathway, hypermethylation-mediated silencing of miRNA-encoding genes, and/or histone modifications, epigenetic silencing through methylation of the promoter regions has emerged as the major mechanism of silencing/downregulation of tumor suppressor miRNAs [12,13]. Also, the two major risk factors of oral cancer, namely smoking and alcohol consumption are reported to impact the epigenetic regulation of various protein-coding genes and miRNAs that are directly involved in OSCC carcinogenesis [14]. Thus, the above observations emphasize a critical role of epigenetically regulated tumor suppressor miRNAs in OSCC pathogenesis.

For the identification of epigenetically regulated miRNAs, a majority of studies employ chromatin-modifying agents or epigenetic modifiers such as DNA hypomethylation agents (DNA methyltransferase inhibitors) like 5-Azacytidine or histone deacetylase (HDAC) inhibitors like 4-phenylbutyric acid (PBA) and trichostatin A (TSA), and in most cases a combination of both to treat cells from cancer cell lines and later adopt a microarray-based approach to identify miRNAs which are differentially expressed between untreated and drug-treated cells [13, 15]. In the present study, by comparing the miRNA expression profiles of 5-Azacytidine and vehicle control-treated OSCC cells followed by the microRNA microarray analysis, we have identified a total of 50 DNA methylation silenced/downregulated miRNAs, and investigated the role of one of these, miR-6741-3p in great details.

## Results

### Analysis of 5-Azacytidine and vehicle control-treated SCC131 cells by miRNA microarray

In order to identify methylation silenced/downregulated tumor suppressor miRNAs, cells from an OSCC cell line SCC131 were treated separately with 5-Azacytidine and vehicle-control (DMSO). To increase sensitivity in the microRNA microarray analysis, total RNA was isolated using a mirVana™ miRNA isolation kit as it specifically enriches for small RNAs (<200 nucleotides). To determine if the 5-Azacytidine treatment of SCC131 cells has worked, the expression of a known tumor suppressor gene *MCPH1*, which is already reported to be epigenetically silenced in SCC131 cells [16] and shows increased expression following 5-Azacytidine treatment, was checked by qRT-PCR (Fig 1A), and the results indicated that it was indeed upregulated following the treatment, suggesting that the treatment of SCC131 cells with 5-Azacytidine has worked. Following this, the expression of 2,549 mature miRNAs was investigated by the microRNA microarray analysis using SurePrint G3 8×60K Human miRNA Microarray chips. Using a cut-off of 0.8-fold expression change, 50 miRNAs were found to be upregulated and 28 miRNAs were found to be downregulated after the 5-Azacytidine treatment compared to the vehicle control (Fig 1B, S1 Table). We decided to work with one of these microRNAs, miR-6741-3p for further analysis as it was found to be upregulated, showed a mean fold expression of 4.96 following the 5-Azacytidine treatment, and there is a lack of reports regarding its role in tumorigenesis (S1 Table).

**Fig 1.**
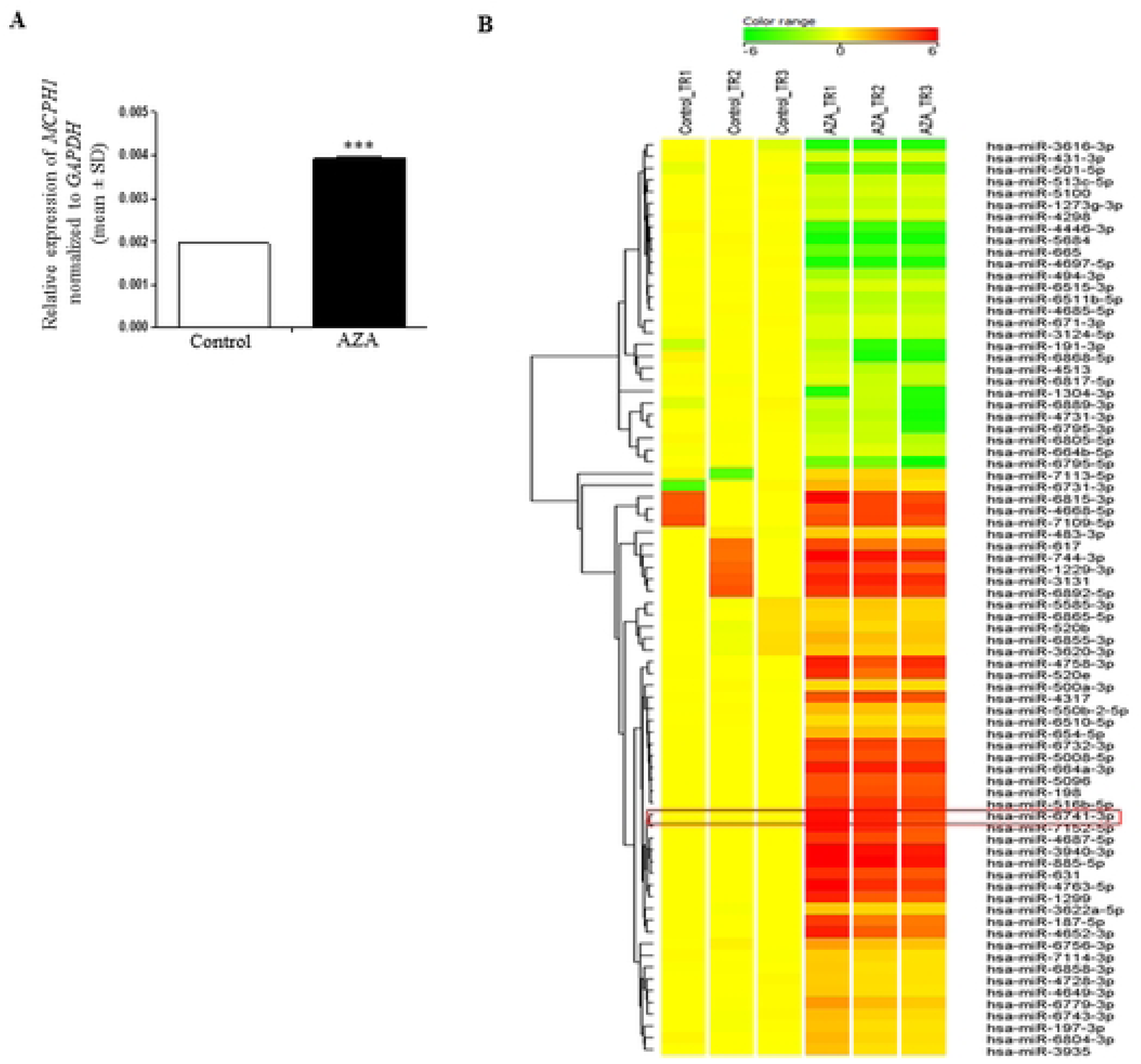
MicroRNA microarray analysis of 5-Azacytidine and control treated SCC131 cells. A) The expression of the tumor suppressor gene *MCPH1* was found to be upregulated following 5-Azacytidine treatment of SCC131 cells by qRT-PCR. B) A heatmap depicting the upregulation of 50 miRNAs and downregulation of 28 miRNAs expression following 5-Azacytidine treatment of SCC131 cells. The change in expression of miR-6741-3p is highlighted. Each bar for qRT-PCR is an average of 2 technical replicates. Control_TR1, Control_TR2 and Control_TR3 represent technical replicates of RNA from DMSO treated cells. AZA_TR1, AZA_TR2 and AZA_TR3 are technical replicates of RNA from 5-Aacytidine treated cells. *Abbreviation*: AZA, 5-Azacytidine.

### Validation of miR-6741-3p upregulation after 5-Azacytidine treatment of SCC131 cells by qRT-PCR

The upregulation of miR-6741-3p following 5-Azacytidine treatment was validated by qRT-PCR using RT6 and short-miR specific primers (S2 Table). As expected, a significant upregulation in miR-6741-3p expression was observed in 5-Azacytidine treated cells compared to the vehicle control treated cells, thus validating the miRNA microarray data (S1 Fig).

### miR-6741-3p overexpression decreases the proliferation of SCC131 and SCC084 cells

Epigenetic silencing due to DNA methylation is one of the key mechanisms for the repression of tumor suppressor miRNAs. As miR-6741-3p was found to be upregulated following 5-Azacytidine treatment of SCC131 cells, we hypothesized that it could be a tumor suppressor miRNA and therefore decided to ascertain its involvement in the regulation of various aspects of cancer cells. To check the involvement of miR-6741-3p in controlling cell proliferation, we transiently transfected SCC131 and SCC084 cells with pcDNA3-*EGFP* (vector control) and pmiR-6741 separately and performed the trypan blue dye exclusion assay. The results showed that miR-6741-3p decreased cell proliferation compared to the vector control in both the cell lines, thus suggesting its role as a tumor suppressor miRNA (S2 Fig).

### *In silico* identification of gene target(s) of miR-6741-3p

Three different mRNA target prediction algorithms (e.g., TargetScan, miRDB, and DIANA microT-CDS) were used to identify gene target(s) for miR-6741-3p. We found *SRSF3* (serine/arginine-rich splicing factor 3), *C6ORF89* (chromosome 6 open reading frame 89), *NDST2* (N-deacetylase/N-sulfotransferase (heparan glucosaminyl) 2), and *MKX* (mohawk homeobox) as potential gene targets of miR-6741-3p predicted by all the three algorithms (S3 Table). However, based on the literature survey, we first decided to check if miR-6741-3p can target *SRSF3* as the involvement of *SRSF3* in the pathogenesis of various cancers including OSCC has already been reported.

### Validation of *SRSF3* as a gene target for miR-6741-3p

To check if *SRSF3* is indeed regulated by miR-6741-3p, we transfected the vector and pmiR-6741 constructs separately in SCC131 cells and assessed the levels of miR-6741-3p and the levels of *SRSF3* transcript and protein (Fig 2A). We observed that miR-6741-3p downregulated the level of SRSF3 protein while the level of *SRSF3* transcript remained unchanged in cells transfected with pmiR-6741 as compared to those transfected with the vector, suggesting that it regulates *SRSF3* expression at the translational level only (Fig 2A). We also transfected different quantities of the pmiR-6741 overexpression construct in SCC131 cells and analysed the expression of *SRSF3* by Western blotting and qRT-PCR. As expected, we observed that miR-6741-3p downregulated the level of *SRSF3* protein in a dose-dependent manner, while the level of *SRSF3* transcript remained unchanged (Fig 2B).

**Fig 2.**
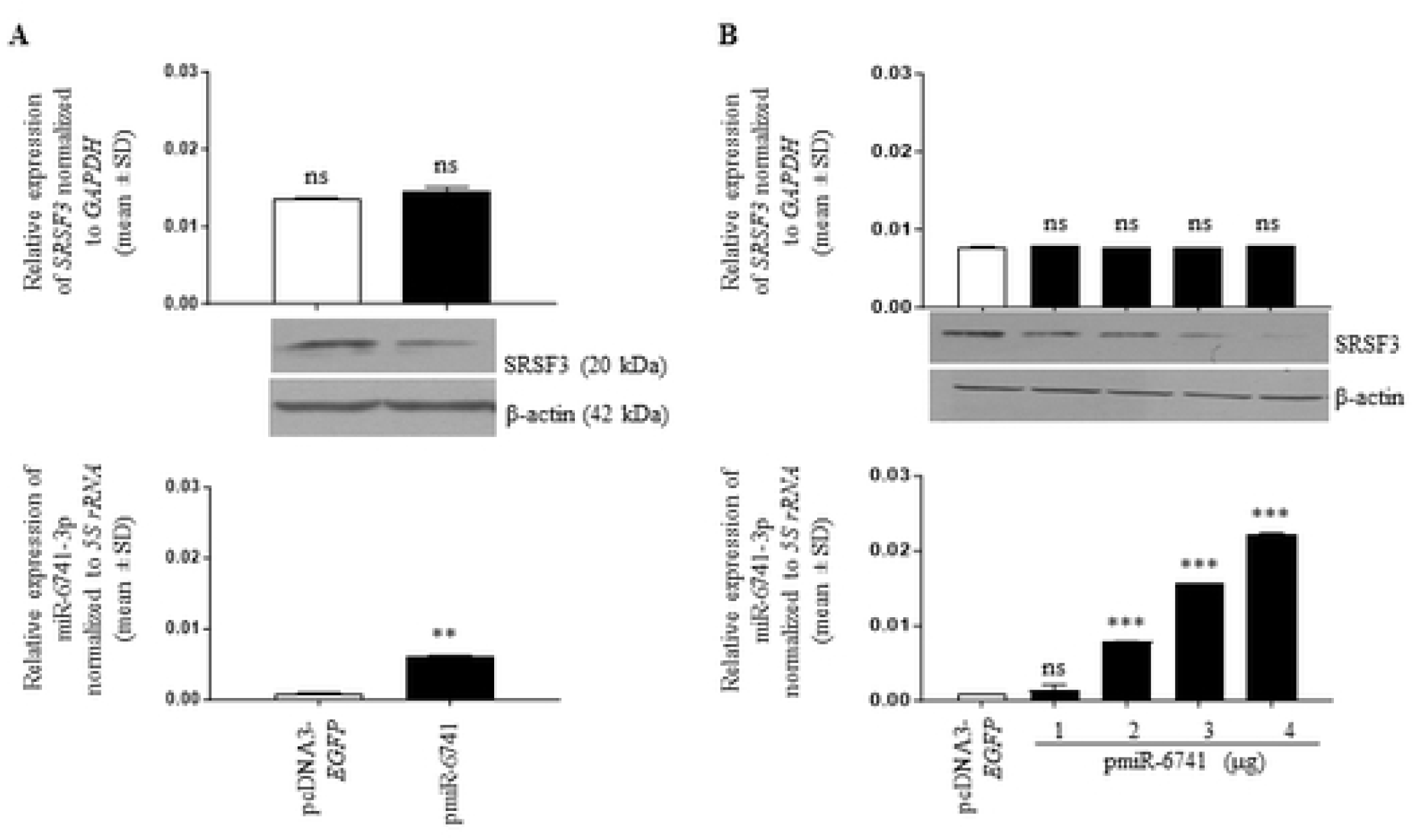
Identification of *SRSF3* as a gene target for miR-6741-3p. A) SRSF3 protein level decreases, while the transcript level remains unchanged on overexpression of miR-6741-3p using the pmiR-6741 construct compared to the vector control. B) A dose-dependent regulation of *SRSF3* at the translational level only is seen on increasing the doses of miR-6741-3p using the pmiR-6741 construct compared to the vector control. For qRT-PCR data, each bar is an average of 2 technical replicates.

### Confirmation of a direct interaction between miR-6741-3p and the 3’UTR of *SRSF3* by dual-luciferase reporter assay

Our bioinformatics analysis predicted one putative target site (TS) for the miR-6741-3p seed region (SR) in the 3’UTR of *SRSF3* from nucleotides 687-694, which is conserved across species (S3A Fig). The dual-luciferase reporter assay was performed to confirm the direct interaction between miR-6741-3p and the 3’UTR of *SRSF3* using different constructs illustrated in S3B Fig. The pMIR-REPORT-*SRSF3*-3’UTR construct with the 3’UTR of *SRSF3* in a sense orientation harbors the TS for miR-6741-3p. The negative control construct, pMIR-REPORT-*SRSF3*-3’UTR-M, is generated by abrogating the TS for miR-6741-3p in the 3’UTR of *SRSF3* by site-directed mutagenesis. To confirm if miR-6741-3p binds directly to the 3’UTR of *SRSF3* and to underscore the importance of the predicted TS, we co-transfected SCC131 cells separately with pMIR-REPORT-*SRSF3*-3’UTR-S and pmiR-6741 or pMIR-REPORT-*SRSF3*-3’UTR-S and the vector pcDNA3-*EGFP* and quantified the luciferase reporter activity. Compared to cells co-transfected with pMIR-REPORT-*SRSF3*-3’UTR-S and pcDNA3-*EGFP*, we observed a significant decrease in luciferase activity in those co-transfected with pMIR-REPORT-*SRSF3*-3’UTR-S and pmiR-6741, confirming that miR-6741-3p binds to the 3’UTR of *SRSF3* in a sequence-specific manner (S3C Fig). Further, as expected, cells co-transfected with pmiR-6741 and pMIR-REPORT-*SRSF3*-3’UTR-Mshowed luciferase activity comparable to those co-transfected with pMIR-REPORT-*SRSF3*-3’UTR-S and pcDNA3-*EGFP* due to the absence of miR-6741-3p TS in the pMIR-REPORT-*SRSF3*-3’UTR-M construct (S3C Fig). These observations suggested that miR-6741-3p binds to the TS in the 3’UTR of *SRSF3* directly in a sequence-specific manner.

### 5-Azacytidine treatment of SCC131 cells upregulates miR-6741-3p expression and downregulates SRSF3

As mentioned earlier, miR-6741-3p was found to be significantly upregulated in 5-Azacytidine-treated SCC131 cells compared to the vehicle control-treated cells (S1 Fig). Since *SRSF3* was identified as the gene target for miR-6741-3p, we next checked the levels of both *SRSF3* transcript and protein in 5-Azacytidine and vehicle control-treated SCC131 cells. As expected, the level of SRSF3 protein was reduced with a concomitant increase in the level of miR-6741-3p in 5-Azacytine treated cells as compared to vehicle control-treated cells (S4 Fig), while no change in the level of *SRSF3* transcript was observed in 5-Azacytidine-treated cells as compared to vehicle control-treated cells (S4 Fig). These observations further underscore the importance of miR-6741-3p-mediated regulation of *SRSF3*.

### Physiological relevance of the interaction between miR-6741-3p and *SRSF3* in cell lines and OSCC patient samples

As mentioned above, the regulation of *SRSF3* by miR-6741-3p is at the translational level. To check the physiological relevance of their interaction, we checked the levels of both miR-6741-3p and SRSF3 protein across seven different cell lines, namely SCC131, SCC084, A549, HeLa, HEK293T, U87, and MCF-7, and in 36 matched normal oral tissue and OSCC samples from patients. In general, an inverse correlation was observed between the expression of miR-6741-3p and SRSF3 across the cell lines that we have tested, indicating that this interaction is of physiological relevance (S5 Fig). For example, the level of miR-6741-3p is highest in U87 cells with almost no expression of SRSF3 protein (S5 Fig). In the case of the OSCC patient samples, miR-6741-3p was found to be significantly downregulated in 16/36 tumor samples (viz., patient no. 3, 8, 33, 46, 49, 56, 64, 2, 5, 6, 10, 17, 31, 45, 48, and 59) as compared to their matched normal oral tissues (S6 Fig, upper panel). Further, we found SRSF3 to be upregulated in 12/36 OSCC samples (viz., patient no. 54, 3, 8, 33, 49, 53, 62, 2, 43, 51, 57, and 66) as compared to matched normal oral tissues (S6 Fig, lower panel). We also found miR-6741-3p to be upregulated in 16/36 OSCC samples (viz., patient no. 54, 68, 47, 52, 53, 55, 14, 32, 43, 44, 50, 51, 60, 61, 65, and 67) as compared to matched normal oral tissues, and no change in its level between normal oral tissue and tumors in 4/36 samples (viz., patient no. 63, 62, 57, and 66) analyzed (S6 Fig, upper panel). SRSF3 was found to be downregulated in 24/36 OSCC samples (viz., patient no. 63, 68, 46, 47, 52, 55, 56, 64, 5, 6, 10, 14, 17, 31, 32, 44, 45, 48, 50, 59, 60, 61, 65, and 67) as compared to matched normal oral tissues (S6 Fig, lower panel). Overall, an inverse correlation was observed between the levels of miR-6741-3p and SRSF3 in 17/36 (47.22%; patient no. 68, 3, 8, 33, 47, 49, 52, 55, 2, 14, 32, 44, 50, 60, 61, 65, and 67) matched OSCC patient samples analyzed (S6 Fig).

### *SRSF3* overexpression increases cell proliferation

To study the role of *SRSF3* in various aspects of cancerous cells like proliferation, apoptosis, and anchorage-independent growth, we generated the p*SRSF3* overexpression construct. Using the same, the effect of *SRSF3* overexpression on proliferation of SCC131 and SCC084 cells was checked using the trypan blue dye exclusion assay. It was observed that *SRSF3* overexpressing cells showed increased cell proliferation compared to those transfected with the vector control in both the cell lines, suggesting that *SRSF3* positively regulates cell proliferation (S7 Fig).

### Expression of SRSF3 depends on the presence or absence of its 3’UTR

To check the effect of miR-6741-3p-mediated regulation of *SRSF3* on its expression and function, we generated two different *SRSF3* constructs by inserting the 3’UTR of *SRSF3* downstream to the *SRSF3*-ORF in the p*SRSF3* construct. The two *SRSF3* constructs are as follows: p*SRSF3*-3’UTR-S with *SRSF3* ORF with its wild-type 3’UTR in a sense orientation and thus harboring a functional TS and p*SRSF3*-3’UTR-M with *SRSF3* ORF with the mutated TS in its 3’UTR in a sense orientation. We then co-transfected both SCC131 and SCC084 cells with pmiR-6741 and different *SRSF3* overexpression constructs or the vector control and performed the Western blot analysis (S8 Fig). The results showed that as compared to cells transfected with the vector control only, cells co-transfected with the vector and pmiR-6741 showed a decreased level of SRSF3 due to the targeting of endogenous SRSF3 by miR-6741-3p (S8 Fig). The level of SRSF3 increased in cells co-transfected with p*SRSF3* and pmiR-6741 as compared to those co-transfected with vector and pmiR-6741 (S8 Fig). The level of SRSF3 was decreased in cells co-transfected with p*SRSF3*-3’UTR-S and pmiR-6741 as compared to those co-transfected with p*SRSF3* and pmiR-6741, because of the presence of a functional TS in the 3’UTR of p*SRSF3*-3’UTR-S. The level of SRSF3 was rescued in cells co-transfected with p*SRSF3*-3’UTR-M and pmiR-6741 as compared to those co-transfected with p*SRSF3*-3’UTR-S and pmiR-6741, due to the presence of a mutated TS in the 3’UTR of p*SRSF3*-3’UTR-M (S8 Fig). These observations suggested that the expression of *SRSF3* depends on the presence or absence of its 3’UTR and is, in part, regulated by miR-6741-3p.

### miR-6741-3p regulates cell proliferation, in part, by targeting the 3’UTR of *SRSF3*

To elucidate the effect of miR-6741-3p-mediated regulation of *SRSF3* on cell proliferation, we co-transfected different *SRSF3* overexpression constructs along with pmiR-6741 or vector control in both SCC131 and SCC084 cells and performed the trypan blue dye exclusion assay. As expected, we observed decreased proliferation of cells co-transfected with vector and pmiR-6741 as compared to those transfected with vector only (Fig 3). Cells co-transfected with pmiR-6741 and p*SRSF3*-3’UTR-S showed decreased cell proliferation as compared to those co-transfected with pmiR-6741 and p*SRSF3*, due to the presence of a functional TS in the 3’UTR of p*SRSF3*-3’UTR-S (Fig 3). As expected, no difference in cell proliferation was observed in cells co-transfected with pmiR-6741 and p*SRSF3* as compared to those co-transfected with pmiR-6741 and p*SRSF3*-3’UTR-M, due to the absence of a functional TS in 3’UTR of p*SRSF3*-3’UTR-M (Fig 3). Similar results were obtained in both cell lines (Fig 3). The above observations indicate that miR-6741-3p negatively regulates cell proliferation, in part, by targeting the 3’UTR of *SRSF3*.

**Fig 3.**
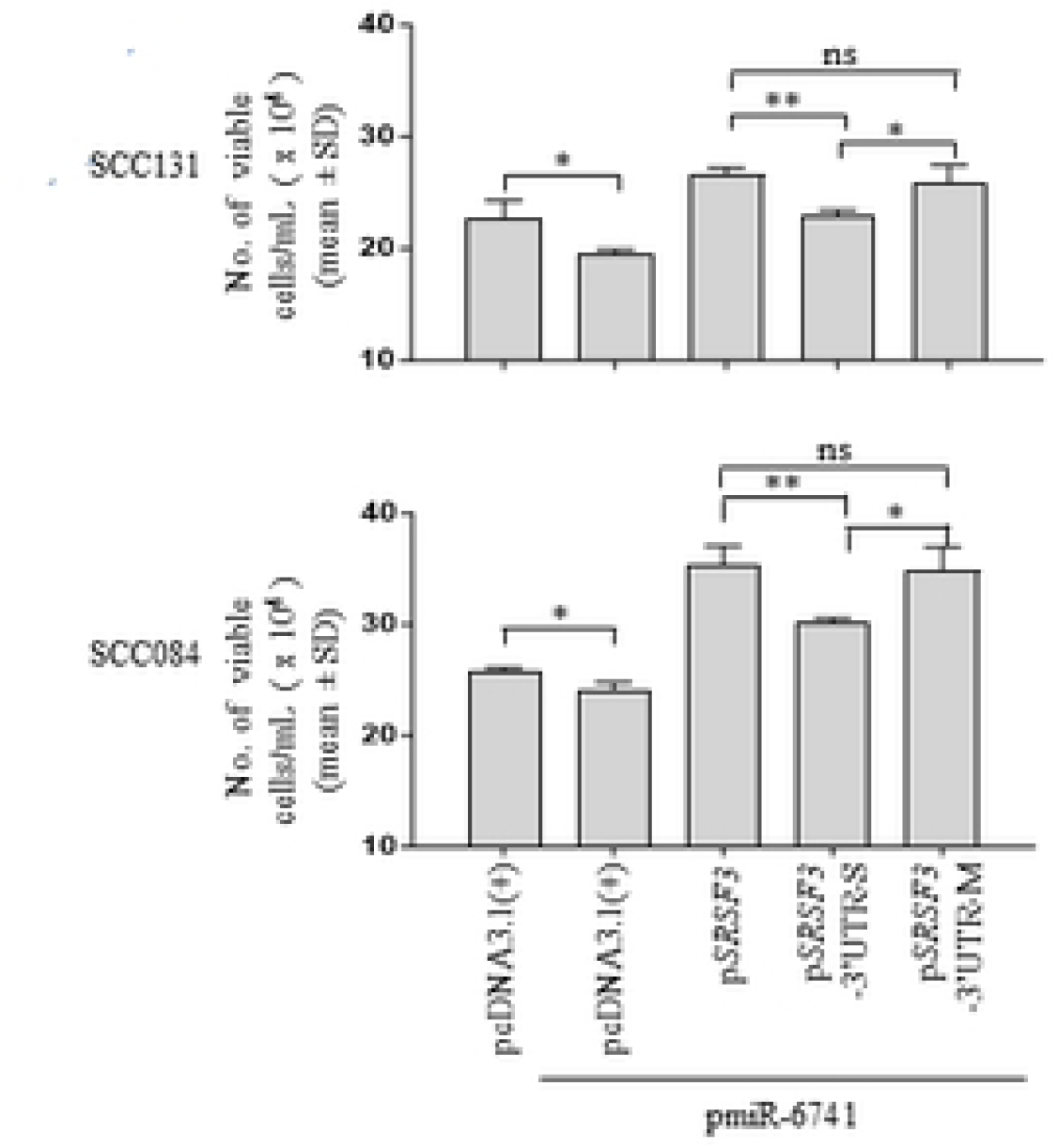
miR-6741-3p regulates cell proliferation, in part, by targeting the 3’UTR of *SRSF3*. Quantitative analysis of cell proliferation by the trypan blue dye exclusion assay in SCC131 and SCC084 cells co-transfected with pmiR-6741 and different *SRSF3* overexpression constructs or vector control.

### miR-6741-3p regulates anchorage-independent growth, in part, by targeting the 3’UTR of *SRSF3*

To analyse the effect of miR-6741-3p-mediated regulation of *SRSF3* on anchorage-independent growth capabilities of cells, we co-transfected different *SRSF3* overexpression constructs with pmiR-6741 or vector control in both SCC131 and SCC084 cells and performed the soft agar colony-forming assay. We used microscopic examination to score for visible colonies at the end of the experiment. As expected, we observed a sharp decrease in the number of colonies in cells co-transfected with pmiR-6741 and the vector as compared to those transfected with vector only (Fig 4). Further, cells co-transfected with pmiR-6741 and p*SRSF3*-3’UTR-S construct harboring a functional TS for miR-6741-3p showed a decrease in the number of colonies compared to those co-transfected with p*SRSF3* and pmiR-6741 (Fig 4). As expected, no significant difference in the number of colonies was observed in cells co-transfected with pmiR-6741 and p*SRSF3* as compared to those co-transfected with pmiR-6741 and p*SRSF3*-3’UTR-M construct harboring a non-functional miR-6741-3p TS (Fig 4). These observations clearly suggest that miR-6741-3p negatively regulates anchorage-independent growth, in part, by targeting the 3’UTR of *SRSF3*.

**Fig 4.**
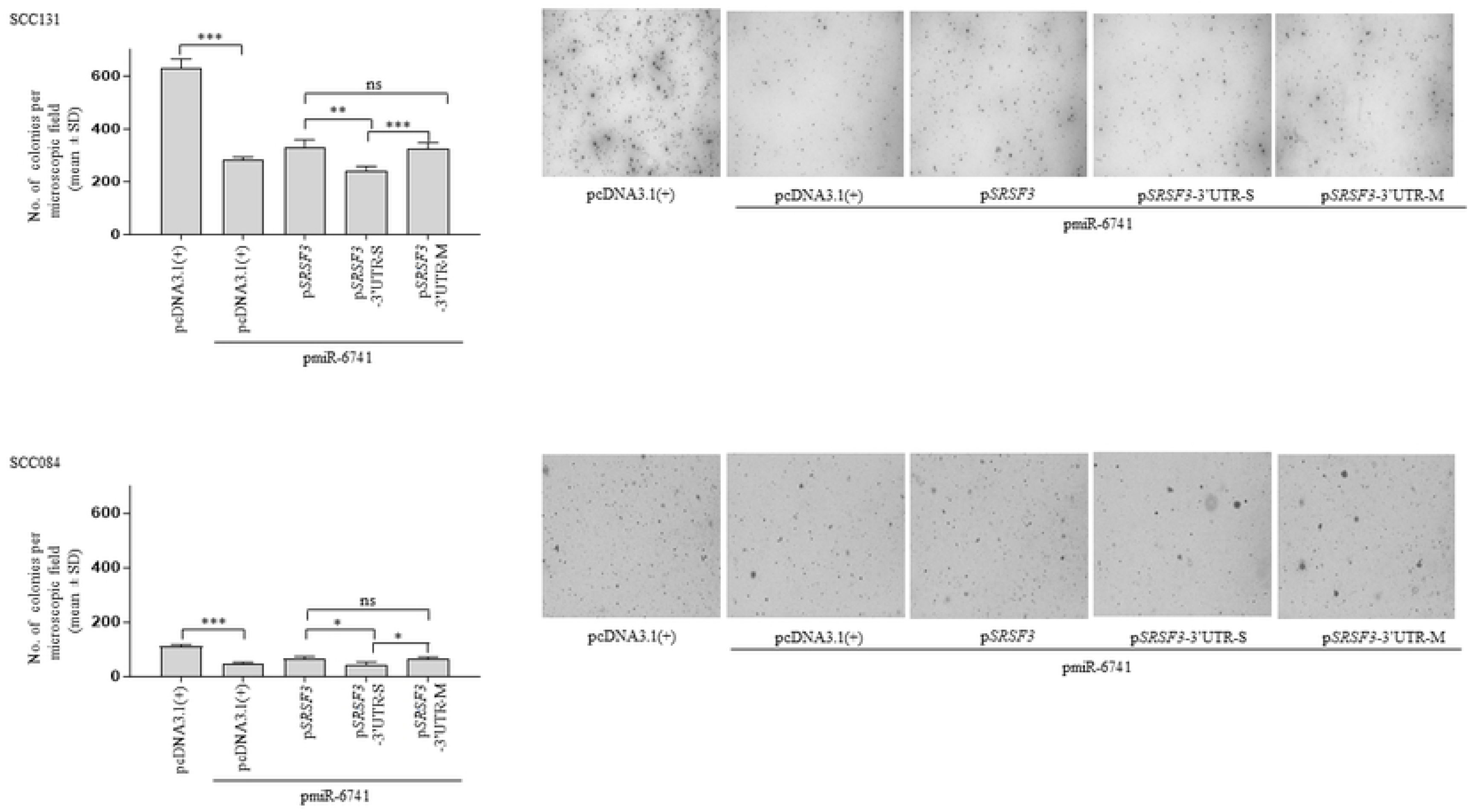
miR-6741-3p regulates anchorage-independent growth, in part, by targeting the 3’UTR of *SRSF3*. Quantitative assessment of the anchorage-independent growth capabilities and representative microphotographs of the colonies for SCC131 (upper panel) and SCC084 (lower panel) cells co-transfected with pmiR-6741 and different *SRSF3* overexpression constructs or vector control by the soft agar colony-forming assay. Each bar is an average of 4 biological replicates.

### miR-6741-3p induces cellular apoptosis independent of *SRSF3*

Using different *SRSF3* overexpression constructs along with pmiR-6741 or vector control, we also analysed the effect of miR-6741-3p-mediated regulation of *SRSF3* on cellular apoptosis in both SCC131 and SCC084 cells. We observed that in both the cell lines, co-transfection of pmiR-6741 with the vector or any of the *SRSF3* constructs led to a significant increase in apoptosis as indicated by an increase in the percentage of Caspase-3 positive cells compared to only vector-transfected cells (Fig 5), suggesting that miR-6741-3p positively regulates apoptosis. However, no difference in the rate of apoptosis was found among cells transfected with any of the *SRSF3* constructs and pmiR-6741 (Fig 5), suggesting that *SRSF3* has no effect on apoptosis. This was further confirmed by transfecting the vector or the p*SRSF3* construct separately in cells from both the cell lines and assessing the rate of apoptosis. The results showed no change in the rate of apoptosis between vector control and p*SRSF3* transfected cells (Fig 5). These observations suggest that miR-6741-3p induces cellular apoptosis in both SCC131 and SCC084 cells independent of *SRSF3*.

**Fig 5.**
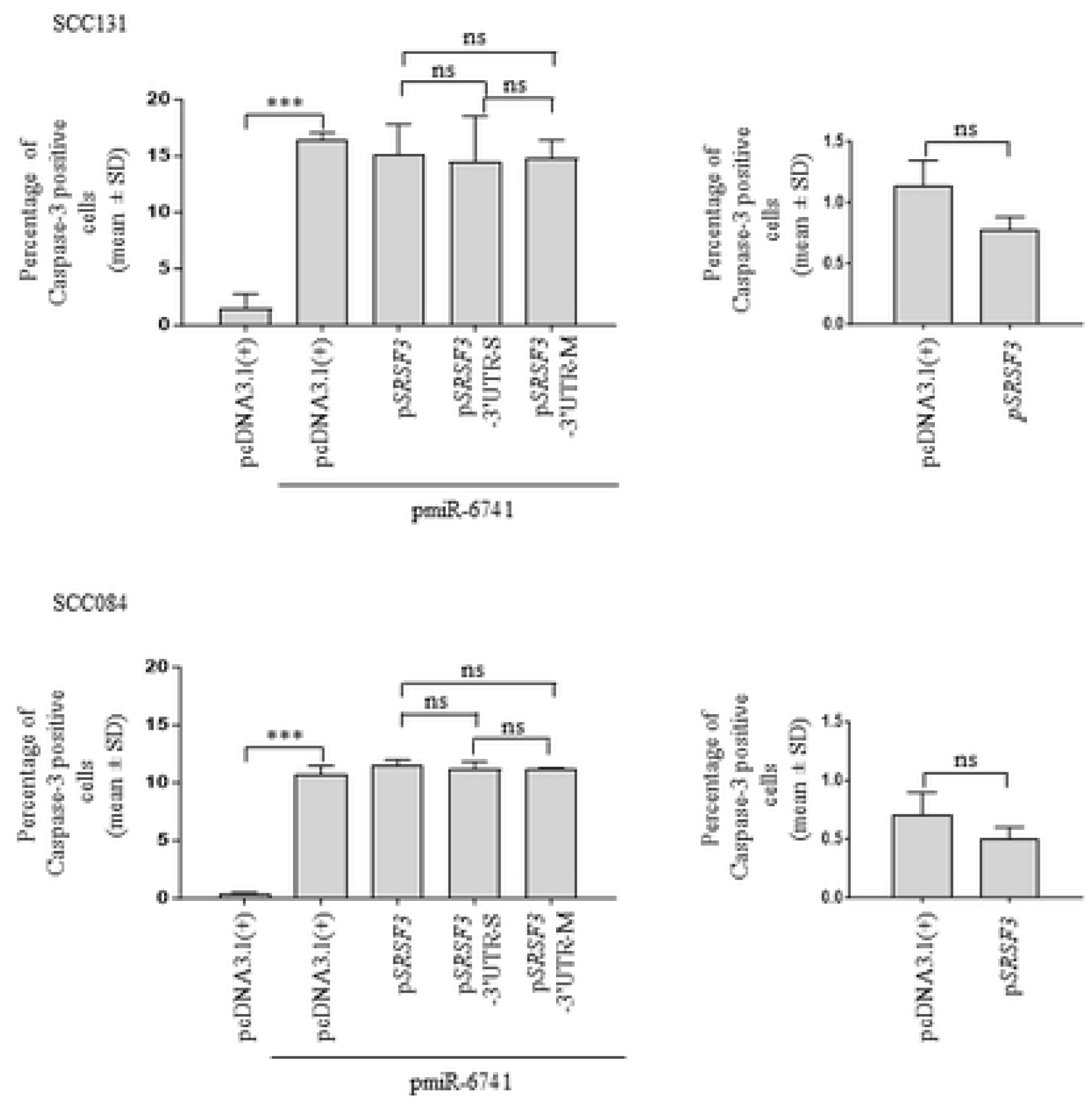
miR-6741-3p induces apoptosis independent of *SRSF3*. Quantitative analysis of the rate of apoptosis as assessed by the percentage of Caspase-3 positive cells in SCC131 (upper panel) and SCC084 (lower panel) cells co-transfected with pmiR-6741 and different *SRSF3* overexpression constructs or vector control. Note, there is no change in the rate of apoptosis on overexpression of *SRSF3* compared to the vector control in both SCC131 and SCC084 cells. Each bar is an average of 3 biological replicates.

### Optimization of dosage for miR-6741-3p mimic and inhibitor in SCC131 cells

We wanted to explore the potential of a synthetic miR-6741-3p mimic and an inhibitor in regulating the levels of SRSF3. To this end, we transfected SCC131 cells with different quantities of mimic and inhibitor for optimization of the dosage. The results showed that 1,500 nM of miR-6741-3p mimic was sufficient to decrease the level of SRSF3 in SCC131 cells (S9A Fig). In the case of miR-6741-3p inhibitor, both 2,000 nM and 3,000 nM dosages were found to be effective in increasing the level of SRSF3 in SCC131 cells in a dose-dependent manner (S9B Fig). As expected, the qRT-PCR analysis showed an increased level of miR-6741-3p in mimic-treated cells and its decreased level in inhibitor-treated cells compared to those treated with controls, confirming their specificity (S9 Fig).

### Restoration of miR-6741-3p by a mimic suppresses *in vivo* tumor growth, while its inhibition by an inhibitor promotes *in vivo* tumor growth in nude mice

Our *in vitro* studies hinted toward an anti-tumor activity of miR-6741-3p. Based on these observations, we hypothesized that the restoration of miR-6741-3p level by a synthetic miR-6741-3p mimic and, in turn, reducing the levels of SRSF3 in OSCC cells might have an anti-tumor effect *in vivo*. Conversely, decreasing the level of miR-6741-3p by a synthetic miR-6741-3p inhibitor and, in turn, increasing the level of SRSF3 in OSCC cells might promote tumor formation *in vivo*. We, therefore, decided to test this hypothesis using *in vivo* pre-treated OSCC xenograft nude mouse model. To this end, we injected equal numbers of SCC131 cells that were pre-transfected with 1,500 nM of miR-6741-3p mimic or 1,500 nM of mimic control separately into the right flanks of female nude mice. In another experimental set, we injected equal numbers of SCC131 cells that were pre-transfected with 3,000 nM of miR-6741-3p inhibitor or 3,000 nM of inhibitor control separately into the left flanks of the female nude mice. The mice were monitored for OSCC xenograft (tumor) growth until 26 days for the mimic group and 29 days for the inhibitor group. As expected, nude mouse xenografts with miR-6741-3p mimic had significantly reduced volumes in comparison to those with control (S10A and S10B Figs). Also, the tumor weights were reduced in mice treated with miR-6741-3p mimic compared to those treated with mimic control; however, the difference was not statistically significant (S10C Fig). Similarly, as expected, nude mouse xenografts with miR-6741-3p inhibitor had significantly increased tumor volumes in comparison to those with inhibitor control (S10D Fig and S10E). As expected, the tumor weights were increased in mice treated with miR-6741-3p inhibitor compared to those treated with inhibitor control; however, the difference was not statistically significant (S10F Fig). Taken together, these observations suggest that miR-6741-3p inhibits tumor growth *in vivo*, in part, by targeting the 3’UTR of *SRSF3*.

### Overexpression of miR-6741-3p leads to a decrease in activation, while overexpression of *SRSF3* leads to activation of the PI3K/AKT/MTOR and ERK/MAP pathways

The PI3K-AKT-MTOR pathway, a central hub for controlling cellular proliferation and growth, is the most frequently activated pathway in OSCC. Apart from this, the ERK/MAPK pathway is also frequently deregulated in various cancers including OSCC. We, therefore, decided to analyze the activation of these two critical pathways on overexpression of miR-6741-3p or *SRSF3*, using the Western blot analysis. As read-outs for the activated PI3K-AKT-MTOR pathway, we decided to check the levels of phospho– and total-S6K1. Similarly, we checked the levels of phospho– and total-ERK1/2 as read-outs for the activated ERK/MAPK signaling pathway. The results showed decreased levels of phospho– and total-S6K1 as well as phospho– and total-ERK1/2 levels in both SCC131 and SCC084 cells, following overexpression of miR-6741-3p (S11A Fig). As expected, overexpression of miR-6741-3p led to a decreased level of SRSF3 in both SCC131 and SCC084 cells (S11A Fig). Overexpression of *SRSF3* on the other hand led to increased levels of phospho– and total-S6K1 as well as phospho– and total-ERK1/2 levels in both SCC131 and SCC084 cells (S11B Fig). The above observations suggest that miR-6741-3p decreases signaling through both the PI3K-AKT-MTOR and the ERK/MAPK pathways, in part, by regulating SRSF3, and there seems to be a miR-6741-3p-SRSF3-ERK1/2-S6K1 axis.

### Putative *MIR6741* promoter lacks any promoter activity

We wanted to identify the mechanism for the upregulation of miR-6741-3p following the 5-Azacytidine treatment of SCC131 cells. We hypothesized that miR-6741-3p is upregulated following 5-Azacytidine treatment due to demethylation of its promoter.

To test this hypothesis, the putative *MIR6741* promoter sequence encompassing the proximal region of the *MIR6741* locus was retrieved from the DBTSS database [17] (S12A Fig) and cloned in the pGL3-Basic vector, a promoterless vector. To check if additional regulatory sequences are needed for the promoter to function, we generated another construct (pmiR-6741-F2) by cloning a larger fragment that encompassed the predicted putative promoter sequence along with additional upstream and downstream sequences (S12B Fig). The two constructs for the putative *MIR6741* promoter are schematically represented in S12C Fig. The two putative promoter constructs (pmiR-6741-F1 and pmiR-6741-F2) along with the control pGL3-Control and pGL3-Basic were then transfected separately in SCC131 cells and the dual-luciferase reporter assay was performed. However, the results showed no promoter activity for both the putative *MIR6741* promoter constructs (S13 Fig), indicating that the proximal region of the *MIR6741* locus does not represent the *MIR6741* promoter.

## Discussion

The present study was focused on the identification of novel tumor suppressor miRNAs involved in OSCC pathogenesis at a genome-wide scale. The microRNA microarray analysis of cells from an OSCC cell line SCC131 treated with 5-Azacytidine and vehicle control (DMSO) led to the identification of 50 upregulated and 28 downregulated miRNAs (Fig 1B,S1 Table). miR-6741-3p was one of the 50 upregulated miRNAs (tumor suppressors) that was validated by qRT-PCR (Figs 1B and S1) and studied in detail. It is a poorly conserved intronic miRNA present in the intron between exons 3 and 4 of the transcript variant I of the host gene *PYCR2* (Pyrroline-5-carboxylate reductase family, member2) and was discovered recently by Ladewig and co-workers [18]. The physiological function of this miRNA has not been annotated and there are very few reports of its involvement in disease conditions [19,20].

Like the classical tumor suppressor genes, one of the defining characteristics of a tumor suppressor miRNA is its ability to suppress the proliferation of cancer cells. In our study, the overexpression of miR-6741-3p in SCC131 and SCC084 cells decreased cell proliferation, confirming its tumor-suppressive nature (S2 Fig). Tumor suppressor miRNAs exert their effect through the repression of their target oncogenic mRNA networks, leading to an inhibition of tumorigenesis [21]. Using bioinformatics analysis, the oncogene *SRSF3* was identified as a potential target for miR-6741-3p (S3 Table). In this study, we showed for the first time that the overexpression of miR-6741-3p downregulated *SRSF3* at the protein level in OSCC cells, but there was no change at the RNA level, indicating that translational inhibition and not mRNA degradation is involved in miR-6741-3p-mediated suppression of *SRSF3* (Fig 2). Using a combination of computational prediction and dual-luciferase reporter assay, we established that the oncogene *SRSF3* is indeed an evolutionarily conserved direct target for miR-6741-3p (S3 Fig).

SRSF3 is a multifunctional protein belonging to the SR family of proteins and apart from its classical role in regulating constitutive and alternative splicing, it also regulates several cellular processes like mRNA export, alternative polyadenylation, miRNA biogenesis, transcription termination, DNA repair, nuclear RNA quality control, stress granule assembly, maintenance of transcriptome integrity of developing oocytes and regulation of pluripotency [22]. It promotes tumorigenesis by regulating the expression of a plethora of protein-coding genes as well as miRNAs that are directly involved in carcinogenesis [23–26]. It is overexpressed in a wide variety of cancers like cancers of the lungs, skin, stomach, liver, cervix, bladder, breasts, colon, kidneys, thyroid, ovaries, various mesenchymal tissues as well as OSCC [27–29]. Gene amplification and impairment of *SRSF3* autoregulation have been attributed to its overexpression in at least a subset of these cancers [27,30]. As we identified *SRSF3* as a direct target for miR-6741-3p, we proposed that the downregulation of miR-6741-3p might also be responsible for the increased expression of SRSF3 in OSCC and other cancers and might play a critical role in tumorigenesis. Our finding of an inverse correlation between the expression of miR-6741-3p and SRSF3 in various cell lines (S5 Fig) and a subset (17/36; 47.22%) of paired normal oral tissue and OSCC samples (S6 Fig) supported that indeed miR-6741-3p-mediated regulation of *SRSF3* is of physiological and biological relevance. However, we were not able to observe any correlation in 19/36 (52.78%) OSCC patient samples. The lack of correlation in the levels of miR-6741-3p and SRSF3 in these OSCC samples could be attributed to factors like tumor heterogeneity, splicing factors redundancy, involvement of additional levels of regulation of *SRSF3*, variable etiopathogenesis, and heterogeneous genetic constitution of each patient [31]. Similar discrepancies in the expression of miRNAs and their target genes across different studies have been reported earlier. For example, Mallela et al. [32] found an inverse correlation in the levels of miR-130a and its target gene *TSC1* in 19/36 (52.78%) OSCC samples only. Rather et al. [33] observed an inverse correlation in the levels of miR-155 and its target gene *CDC73* in 10/18 (55.56%) OSCC samples only.

Given the fact that SRSF3 is overexpressed in multiple cancers, including OSCC and acts as an oncogene, we decided to investigate how miR-6741-3p-mediated regulation of *SRSF3* affects its oncogenic function in OSCC cells *in vitro* and *in vivo*. To assess the same, we incorporated the sense and mutant 3’UTR of *SRSF3* downstream to the *SRSF3* ORF in the p*SRSF3* construct and co-transfected these constructs (p*SRSF3*-3’UTR-S and p*SRSF3*-3’UTR-M) separately in OSCC cells with pmiR-6741. This approach helped us to concurrently confirm that the change in expression of *SRSF3* is due to the interaction between its 3’UTR and miR-6741-3p and analyzed the effect of this interaction on the oncogenic function of SRSF3. The Western blot analysis revealed that the presence of the wild-type 3’UTR and not the mutant 3’UTR in the expression vector dramatically inhibited the production of SRSF3 protein (S8 Fig) in miR-6741-overexpressing OSCC cells, indicating that the expression of *SRSF3* is modulated by the interaction of miR-6741-3p with its 3’UTR. We then co-transfected all the *SRSF3* constructs (p*SRSF3*, p*SRSF3*-3’UTR-S, and p*SRSF3*-3’UTR-M) along with the pmiR-6741 construct and investigated if the miR-6741-3p-mediated knockdown of *SRSF3* is reflected on cell proliferation, anchorage-independent growth, and apoptosis of OSCC cells (Figs 3-5). In the presence of the pmiR-6741 construct, the overexpression of SRSF3 protein using the constructs without its 3’UTR (p*SRSF3*) or with the mutated 3’UTR (p*SRSF3*-3’UTR-M) promoted the proliferation and anchorage-independent growth of OSCC cells (Figs 3 and 4). This is in line with other studies where the overexpression of SRSF3 promoted proliferation and anchorage-independent growth of cancer cells, while the knockdown of *SRSF3* suppressed proliferation and anchorage-independent growth [27,28,34]. However, contrary to the earlier studies [27,28,35–36] which demonstrated the anti-apoptotic property of SRSF3, in our study, overexpression of SRSF3 protein using any of the three constructs was not able to rescue miR-6741-3p-induced apoptosis, indicating that SRSF3 has no effect on apoptosis of OSCC cells (Fig 5). Taken together, the *in vitro* studies demonstrated that miR-6741-3p suppresses proliferation and anchorage-independent growth of OSCC cells, in part, by targeting the 3’UTR of *SRSF3*, and promotes apoptosis of OSCC cells independent of *SRSF3*.

We have further confirmed the tumor-suppressive properties of miR-6741-3p in oral cancer using an *in vivo* pre-treatment OSCC xenograft nude mice model system. In our study, we demonstrated that restoration of miR-6741-3p by a mimic suppresses *in vivo* tumor growth, while its inhibition by an inhibitor promotes *in vivo* tumor growth in nude mice. (S10 Fig). Though the observed differences in tumor volume between miR-6741-3p mimic and mimic control-treated group, as well as, miR-6741-3p inhibitor and inhibitor control-treated groups were reflected in tumor weights, the difference was not statistically significant (S10C and S10F Figs). This limitation of the present study can be attributed to inter-animal variation in tumor weights, which in turn was largely due to variation in time required for tumor induction in our animal cohort.

Lastly, in a bid to identify the molecular effectors and pathways affected by miR-6741-3p-mediated regulation of *SRSF3* in OSCC, we focused on two critical pathways namely, PI3K-AKT-MTOR and ERK/MAPK which are frequently deregulated in OSCC and other cancers [37–39]. Our results demonstrated that there is a miR-6741-3p-SRSF3-ERK1/2-S6K1 axis through which miR-6741-3p decreases signaling, in part, by modulating SRSF3 (S11 and S14 Figs). Taken together, our findings from the *in vitro* and *in vivo* assays not only highlight the oncogenic role of SRSF3 during oral carcinogenesis, but also strongly attest to the tumor-suppressive role of miR-6741-3p in OSCC, in part, by targeting *SRSF3*.

Early reports suggested that transcription of intronic miRNAs is linked to the host gene transcription and requires RNA Pol II and splicing machinery for their biogenesis [40,41]. However, Monteys et al. [42] predicted that ∼35% of intronic miRNAs can be transcribed from independent promoters by Pol II or Pol III. The increase in expression of miR-6741-3p following 5-Azacytidine treatment of SCC131 cells (Figs 1B and S1) therefore could be attributed to the demethylation at the *MIR6741* promoter if *MIR6741* has its independent promoter. In our study, *MIR6741* promoter constructs (pmiR-6741-F1 and pmiR-6741-3p-F2) that we generated based on the prediction by the DBTSS database were not able to drive the expression of the luciferase reporter (Figs S12 and S13), indicating that *MIR6741* lacks an independent promoter. We will be thus exploring the underlying mechanism of miR-6741-3p upregulation following 5-Azacytidine treatment of SCC131 cells in the future.

In summary, our study identified for the first time a total of 50 potential tumor suppressor miRNAs in OSCC on a genome-wide scale. The current study clearly demonstrated that the oncogene *SRSF3* is a target for the tumor suppressor miR-6741-3p. We substantiate this conclusion with a combination of *in silico*, *in vitro,* and *in vivo* assays. Further, we suggest that the restoration of miR-6741-3p level by using a synthetic miR-6741-3p mimic could be a potent strategy to treat OSCC and perhaps other cancers.

## Materials and methods

### Cell lines

UPCI: SCC131 (SCC131) and UPCI: SCC084 (SCC084) cell lines are a kind gift from Dr. Susanne M. Gollin (University of Pittsburgh, Pittsburgh, PA, USA). HeLa, A549 and HEK293T cells were procured from the National Centre for Cell Sciences, Pune, India. U87 and MCF-7 cell lines were obtained from Prof. P. Kondaiah’s laboratory, Department of Developmental Biology and Genetics, IISc, Bengaluru, India.

### 5-Azacytidine treatment and miRNA microarray analysis

SCC131 cells were treated separately with 5 µM 5-Azacytidine for 5 days (cat# A1287; Sigma-Aldrich, St. Louis, MO, USA) and vehicle control DMSO (cat# D4540; Sigma-Aldrich, St. Louis, MO, USA), following a standardized laboratory protocol. Following this, the expression of 2,459 mature miRNAs was investigated by the microRNA microarray analysis using SurePrint G3 8×60K Human miRNA Microarray chips (AMADID 70156; Agilent Technologies, Santa Clara, CA, USA).

### *In silico* identification of targets for miR-6741-3p

Three target prediction programs, namely miRDB [43], DIANA-microT-CDS [44,45] and TargetScan [46] were used to identify target genes for miR-6741-3p (S3 Table).

### Sample collection

A total of 36 matched normal oral tissue and OSCC patient samples were ascertained at the Kidwai Memorial Institute of Oncology (KMIO), Bengaluru, from 13^th^ July, 2018 to 14^th^ November, 2018. The study was performed with written informed consent from the patients following approvals from the ethics committee of Kidwai Memorial Institute of Oncology, Bengaluru (approval # KMIO/MEC/021/05.January.2018). This study was conducted in accordance with principles of Helsinki declaration. The samples were obtained as surgically resected tissues from oral cancerous lesions and adjacent normal tissues (taken from the farthest margin of surgical resection) in the RNALater™ (Sigma-Aldrich, St. Louis, MO, USA) and transferred to –80°C until further use. The tumors were staged according to the UICC’s (International Union against Cancer) TNM (Tumor, Node, and Metastasis) classification [47]. The details of the clinicopathological parameters obtained from the patients are summarised in S4 Table.

### Total RNA extraction and qRT-PCR

Total microRNA enriched RNA sample for microRNA microarray analysis was isolated using a mirVana™ miRNA isolation kit (cat# AM1561; Ambion, Austin, TX, USA), according to the manufacturer’s protocol. Total RNA from cell lines and tissues was isolated using TRI Reagent™ (Sigma-Aldrich, St. Louis, MO, USA). RNA was quantitated using a NanoDrop™ 1000 spectrophotometer (Thermo Fisher Scientific, Waltham, MA, USA). First-strand cDNA synthesis was done using 1-2 μg of total RNA and a Verso cDNA Synthesis Kit (Thermo Fischer Scientific, Waltham, MA, USA). The expression of miR-6741-3p was determined as suggested by Sharbati-Tehrani et al. [48]. The qRT-PCR analysis was carried out using a DyNAmo ColorFlash SYBR Green qPCR Kit in a StepOnePlus Real-Time PCR System (Thermo Fischer Scientific, Waltham, MA, USA). *GAPDH* and *5S rRNA* were used as normalizing controls. The following equation, ΔCt_gene_ = Ct_gene_-Ct_normalizing control_, was used to calculate the fold change in expression. Ct represents cycle threshold value, and ΔCt represents the gene expression normalized to *GAPDH* or *5S rRNA*. A two-tailed unpaired t-test was performed using the GraphPad PRISM5 software (GraphPad Software Inc., San Diego, CA, USA) to analyze the statistical significance of the difference in mRNA expression. Details of the RT-PCR primers are given in S2 Table.

### *In silico* identification of the putative MIR6741 promoter

The putative promoter sequence for *MIR6741* was retrieved by an *in silico* search using the DBTSS [17] database.

### Plasmid constructs

miR-6741 (pmiR-6741) and *SRSF3* (p*SRSF3*) overexpression constructs were generated in the pcDNA3-*EGFP* and pcDNA3.1(+) vectors respectively, using human genomic DNA or human cDNA as templates as required and gene-specific PCR primers following a standard laboratory procedure (S5 Table). Different restriction enzyme sites were incorporated in forward and reverse primers to facilitate directional cloning.

To generate the pMIR-REPORT-*SRSF3*-3’UTR-S construct containing the 3’UTR of *SRSF3* at the 3’ end of luciferase ORF in the pMIR-REPORT™ vector (Invitrogen, Waltham, MA, USA), fragments were amplified using specific primers and human genomic DNA as a template and cloned in a sense orientation using a standard laboratory method (S5 Table). The pMIR-REPORT-*SRSF3*-3’UTR-M construct containing the mutated target site in *SRSF3* 3’UTR was also generated by site-directed mutagenesis according to Sambrook et al. [49] using specific primers and pMIR-REPORT-*SRSF3*-3’UTR-S as the template (S6 Table). The p*SRSF3*-3’UTR-S and p*SRSF3*-3’UTR-M constructs carrying the wild-type (WT) and mutant (M) 3’UTR of *SRSF3* hooked downstream to the *SRSF3* ORF were generated by sub-cloning the wild-type and mutant *SRSF3* 3’UTR from pMIR-REPORT-*SRSF3*-3’UTR-S and pMIR-REPORT-*SRSF3*-3’UTR-M constructs respectively in the p*SRSF3* construct. Briefly, the *SRSF3*-3’UTR sense and mutant fragments were excised from the pMIR-REPORT-*SRSF3*-3’UTR-S and pMIR-REPORT-*SRSF3-*3’UTR-M constructs respectively by digestion with *Bam* HI and *Eco* RV restriction enzymes (S5 Table). The digested fragments were then ligated and cloned in the p*SRSF3* construct also digested by the same enzymes to ensure directional cloning to generate the p*SRSF3*-3’UTR-S and p*SRSF3*-3’UTR-M constructs.

To generate the pmiR-6741*-*F1 and pmiR-6741*-*F2 constructs used for promoter validation in the pGL3-Basic vector (Promega, Madison, WI, USA), fragments were amplified using specific primers and human genomic DNA as the template and cloned using a standard laboratory method (S5 Table). The details of the sequences used to generate the different *MIR6741* promoter constructs are given in S7 Table.

All the constructs used in the study were validated by restriction enzyme digestion and Sanger sequencing on a 3730×l DNA Analyzer (Thermo Fisher Scientific, Waltham, MA, USA).

### Cell culture, transient transfection, and reporter assays

All the cell lines were maintained in Dulbecco’s modified eagle medium (DMEM) supplemented with 10% (v/v) fetal bovine serum (FBS) and 1X antibiotic-antimycotic solution [DMEM and 1X antibiotic-antimycotic solution from Sigma-Aldrich, St. Louis, MO, USA; FBS from Thermo Fisher Scientific, Waltham, MA, USA)] in a humidified incubator with 5% CO_2_ at 37°C.

For overexpression studies, SCC131 or SCC084 cells were seeded at a density of 2 × 10^6^ cells/well in a 6-well plate and transiently transfected with an appropriate construct or co-transfected with a combination of constructs using the Lipofectamine 2000 transfection reagent (Thermo Fisher Scientific, Waltham, MA, USA), following the manufacturer’s protocol. Post 48 hr of transfection, total RNA and protein were isolated from the cells. The direct interaction between the 3’UTR of the target gene and miRNA as well as the promoter activity of the generated constructs was validated using the dual-luciferase reporter assay. Briefly, 5 × 10^4^ cells/well were transfected with different constructs as mentioned above. The assay was carried out after 48 hr of transfection in SCC131 cells, using the Dual Luciferase® Reporter Assay System (Promega, Madison, WI, USA) and the VICTOR™ X Multilabel Plate Reader (PerkinElmer, Waltham, MA, USA) [33,50]. The transfection efficiency in the dual-luciferase reporter assay was normalized by co-transfecting with the pRL-*TK* control vector [33,50].

### Western blot hybridization

Protein lysates from cell lines were prepared using the CelLytic™ M Cell Lysis Reagent (Sigma-Aldrich, St. Louis, MO, USA), while CelLytic™ MT Mammalian Tissue Lysis Reagent (Sigma-Aldrich, St. Louis, MO, USA) was used to prepare lysates from oral tissue samples. The proteins were resolved on SDS-PAGE and transferred onto a PVDF membrane (Pall Corp., Port Washington, NY, USA) using a locally made conventional semi-dry or wet transfer apparatus (Bio-Rad™, Hercules, CA, USA) as per the requirement. The membrane was blocked using 5% skimmed milk powder (Nestlé India Ltd., Gurgaon, India) in 1X PBS-Tween^®^20 buffer. The signal was visualized using appropriate primary and secondary antibodies and the Immobilon™ Western Chemiluminescent HRP substrate (Merck, Darmstadt, Germany) and developed on an X-ray film. The anti-mouse β-actin (1:10,000 dilution, cat# A5441; Sigma-Aldrich, St. Louis, MO, USA) was used as a loading control. An anti-SRSF3 antibody (1:2000 dilution, cat# ab198291) was purchased from Abcam (Cambridge, MA, USA). Antibodies such as anti-phospho-p44/42 MAPK (Erk1/2) (Thr202/Tyr204) (1:1000 dilution, cat# 9101), anti-p44/42 MAPK (Erk1/2) (1:1000 dilution, cat# 9102), anti-phospho-p70 S6 Kinase (Thr421/Ser424) (1:1000 dilution, cat# 9204) and anti-p70 S6 Kinase (1:1000 dilution, cat# 9202) were purchased from Cell Signaling Technology (Danvers, MA, USA). The anti-rabbit HRP-conjugated secondary antibody (1.5:5000 dilution, cat# HP03) and anti-mouse HRP-conjugated secondary antibody (1:5000 dilution, cat# HP06) were purchased from Bangalore Genei^®^ (Bengaluru, India).

### Cell proliferation assay

The rate of proliferation of SCC131 and SCC084 cells transfected with an appropriate construct or co-transfected with a combination of constructs was assessed by employing a trypan blue dye exclusion assay as described by Karimi et al. [51].

### Apoptosis assay

The rate of cellular apoptosis was measured using a CaspGLOW™ Fluorescein Active Caspase-3 Staining kit (BioVision, Milpitas, CA, USA), according to the manufacturer’s instructions, and as described by Mallella et al. [32].

### Soft agar colony-forming assay

Tumor cells can overcome anoikis to proliferate and form colonies in suspension within a semi-solid medium such as soft agar [52]. The anchorage-independent growth of cells co-transfected with a combination of constructs was analysed by the soft agar colony-forming assay in 35 mm tissue culture dishes, following a standard laboratory protocol [33].

### *In vivo* assay for tumor growth

The tumor-suppressive property of miR-6741-3p was investigated using an *in vivo* nude mice OSCC xenograft model. The effect of miR-6741-3p overexpression using a synthetic miR-6741-3p mimic and a synthetic miR-6741-3p inhibitor on tumor growth was assayed in 4-6 weeks female BALB/c athymic nude mice. Briefly, 2×10^6^ SCC131 cells were transfected with 1,500 nM of mimic control or miR-6741-3p mimic. Post 24 hr of transfection, 2×10^6^ cells were suspended in 150 μL DPBS and then subcutaneously injected into the right posterior flank of each mouse. The same method was used to inject SCC131 cells transfected with 3,000 nM of inhibitor control or miR-6741-3p inhibitor into the left posterior flank of each mouse. Tumors were allowed to grow in animals of all the four experimental sets, and tumor volumes were measured using a Vernier’s caliper every alternate day till the termination of the experiment. Tumor volumes were calculated using the formula: V= L×W^2^×0.5, where L and W represent the length and width of the tumor respectively. The animals were photographed, and the tumor xenografts were harvested at the end of the study. Harvested xenografts were also photographed and weighed. The study was conducted following approval (#Project Proposal No. 766, dated October 08,2020) from the Institute Animals Ethics Committee, Indian Institute of Science, Bengaluru. All mice were maintained on a 12:12 h light/dark cycle in proper cages with sufficient food and water. Animal experiments were performed in accordance with the National Institutes of Health Guide for the Care and Use of Laboratory Animals and ARRIVE guidelines. The miRNA mimics and inhibitors used in the study-miRIDIAN microRNA hsa-miR-6741-3p-Mimic (cat# C-302786-00-0020), miRIDIAN microRNA hsa-miR-6741-3p-Hairpin Inhibitor (cat# IH-302786-01-0020), miRIDIAN microRNA Mimic Negative Control #1 (cat# CN-001000-01-20), and miRIDIAN microRNA Hairpin Inhibitor Negative Control #1 (cat# IN-001005-01-20)-were all purchased from Dharmacon (Lafayette, CO, USA). All experiments were carried out in accordance with relevant guidelines and regulations.

## Statistical analysis

A two-tailed student’s *t-*test was performed using the GraphPad PRISM5 software (GraphPad Software Inc., San Diego) to analyze the statistical significance of the difference between two data sets. Differences with P-value ≤0.05 (*), P-value <0.01 (**), and P-value <0.001 (***) were considered statistically significant, whereas P-value >0.05 was considered as statistically non-significant (ns).

## Acknowledgments

We are grateful to the patients for providing the normal and tumor oral samples. DAM is grateful to the Council of Scientific and Industrial Research, New Delhi, for the student fellowship (#09/079(2609)/2013-EMR-I). This work was funded by a grant (# BT/PR33054/MED/30/2210/2020) from the Department of Biotechnology, New Delhi.

## Supplementary information

**S1 Fig. Validation of miR-6741-3p upregulation following 5-Azacytidine treatment of SCC131 cells by qRT-PCR.**

**S2 Fig. miR-6741-3p decreases the proliferation of SCC131 and SCC084 cells.**

**S3 Fig. Confirmation of a direct interaction between miR-6741-3p and the 3’UTR of *SRSF3* by the dual-luciferase reporter assay in SCC131 cells.**

**S4 Fig. 5-Azacytidine treatment of SCC131 cells upregulates miR-6741-3p expression and downregulates SRSF3.**

**S5 Fig. Expression analysis of miR-6741-3p and SRSF3 in cell lines.**

**S6 Fig. Expression analysis of miR-6741-3p and SRSF3 in OSCC patient samples.**

**S7 Fig. SRSF3 overexpression increases the proliferation of SCC131 and SCC084 cells.**

**S8 Fig. SRSF3 expression depends on the presence or absence of its 3’UTR.**

**S9 Fig. Optimization of dosage for miR-6741-3p mimic and inhibitor in SCC131 cells**

**S10 Fig. The effect of synthetic miR-6741-3p mimic and inhibitor on SCC131 cell-derived xenografts in nude mice.**

**S11 Fig. MiR-6741-3p decreases signaling through both PI3K-AKT-MTOR and ERK/MAPK pathways, in part, by regulating SRSF3.**

**S12 Fig. Putative promoter sequence for intronic *MIR6741***.

**S13 Fig. The dual-luciferase reporter assay for putative *MIR6741* promoter fragments in SCC131 cells.**

**S14 Fig. Effect of miR-6741-3p-mediated regulation of SRSF3 on the PI3K/AKT/MTOR and the ERK/MAPK pathways.**

**S1 Table. Differentially expressed microRNAs identified by microRNA microarray analysis of 5-Azacytidine and control-treated SCC131 cells.**

**S2 Table. Details of primers used in qRT-PCR.**

**S3 Table. A list of predicted gene targets^ for miR-6741-3p.**

**S4 Table. Clinicopathological parameters of the patients included in the study.**

**S5 Table. Details of plasmid constructs used in the study.**

**S6 Table. Details of constructs generated by site-directed mutagenesis in the study.**

**S7 Table. Details of the putative *MIR6741* promoter fragments cloned in the pGL3-Basic vector.**

**S1_raw_images**

## Notes

### Competing Interest Statement

The authors have declared no competing interest.

